# Lipid A counteracts doxorubicin-induced systemic dysfunction by boosting mitochondrial activity

**DOI:** 10.64898/2026.04.16.719094

**Authors:** Yuga Nakaguma, Yuri Kato, Yara Atef, Tomoya Ito, Akiyuki Nishimura, Motonari Uesugi, Yasunari Kanda, Jun Kunisawa, Motohiro Nishida

**Author notes:** Correspondence and requests for materials should be addressed to Motohiro Nishida, Ph.D., Department of Physiology, Graduate School of Pharmaceutical Sciences, Kyushu University, 3-1-1 Maidashi, Higashi-ku, Fukuoka 812-8582, Japan.

## Abstract

Vaccine adjuvants are critical for enhancing immune responses and sustaining antibody production. Although their safety profiles are well established, assessments have largely focused on metabolic and excretory organs such as the liver and kidneys, with limited attention to the heart. Here, we systematically evaluated the cardiac effects of five representative adjuvants in mice: alum, MF59, AS03, Sigma Adjuvant Systems, and lipid A. None of the adjuvants impaired baseline cardiac contractile function. Notably, lipid A uniquely enhanced mitochondrial respiratory capacity in rat and human induced pluripotent stem cell-derived cardiomyocytes and promoted mitochondrial membrane hyperpolarization. We next examined its therapeutic potential in a doxorubicin (Dox)-induced heart failure model characterized by mitochondrial dysfunction. Co-administration of lipid A with influenza hemagglutinin (HA) antigen significantly ameliorated cardiac dysfunction. In parallel, lipid A prevented the Dox-induced decline in anti-HA antibody titers, an effect associated with preservation of splenic B cell populations. Collectively, these findings reveal a previously unappreciated cytoprotective dimension of lipid A, demonstrating that it not only potentiates immune responses but also counteracts chemotherapy-induced functional decline by enhancing mitochondrial activity.

## Introduction

Vaccine adjuvants enhance the efficacy of vaccines by boosting their immunogenicity (1). Adjuvanted vaccines offer several benefits, including increased antibody titers and improved durability, and have become an important means of disease prevention not only for immunocompromised individuals with underlying conditions but also for the general public (2). A wide variety of adjuvants, each with unique properties, enhance immune responses (3). Alum, one of the most widely used adjuvants, is composed of aluminum hydroxide and promotes a Th2-type immune response, leading to increased production of IgG1 and IgE antibodies (4). Emulsion-based adjuvants such as MF59 and AS03 enhance antigen uptake by antigen-presenting cells and induce robust immune responses through the production of cytokines and chemokines. These adjuvants predominantly promote Th2 responses while also inducing moderate Th1 responses (5, 6). Sigma Adjuvant Systems (SAS) is also an oil-in-water emulsion adjuvant. Other adjuvants that target TLRs also exist. Monophosphorylated lipopolysaccharide (MPL), isolated from LPS, activates TLR4 on antigen-presenting cells (APCs) and induces cytokine release. Subsequently, it induces a potent Th1 response. Furthermore, lipopolysaccharide A (lipid A), a lipopolysaccharide derived from *Alcaligenes faecalis*, possesses adjuvant activity without causing excessive inflammation (7, 8).

In the safety evaluation of such adjuvants, the focus has traditionally been on assessing renal and hepatic toxicity, inflammatory responses, and cytotoxicity (9, 10). However, rare cases of myocarditis have been reported following COVID-19 mRNA vaccination (11, 12), highlighting the importance of carefully evaluating potential cardiac effects as part of vaccine safety assessment. Although adjuvants used in clinical settings have undergone safety assessment, their organ-specific effects, particularly on the heart, have not been fully characterized. Therefore, further investigation is needed.

The heart requires enormous amounts of energy to maintain its beating, and ATP production is carried out by mitochondria. Cardiomyocytes contain a high density of mitochondria, which account for approximately 30% of their volume (13). Mitochondria maintain their function through repeated fusion and fission (14). It has been reported that mitochondrial function and morphology are impaired and cardiac function is suppressed in cases of myocardial infarction, diabetes, and during treatment with some anticancer drugs (15–17). Excessive reactive oxygen species (ROS) produced under pathological conditions damage the mitochondrial electron transport chain and mtDNA, impairing energy production (18). Furthermore, as oxidative stress increases, mitochondria undergo hyperfission and become dysfunctional, leading to heart failure (19). Thus, cardiac mitochondria are a suitable target for evaluating cardiotoxicity.

Doxorubicin (Dox), a well-known cardiotoxic anticancer agent, induces mitochondrial dysfunction in a dose-dependent manner (20). Multiple mechanisms have been reported for Dox-induced cardiotoxicity (21–23). Dox-induced cardiomyopathy is widely used as an experimental model, making it a suitable system to evaluate whether adjuvants induce similar pathological changes (24, 25). Furthermore, patients undergoing anticancer therapy are often immunocompromised, and vaccination is important for infection prevention (26–29). Thus, it is also important to understand how adjuvants affect the immunosuppression caused by anticancer drugs.

In this study, we evaluated the safety of various adjuvants from a novel perspective by focusing on cardiac mitochondrial energy metabolism. Using a Dox-induced heart failure mouse model, we investigated the effects of adjuvants on mitochondrial function and cardiac phenotype.

## Methods

### Material

Alum (Alhydrogel 2%, Cat# vac-alu-10), MPL (MPLA-SM, Cat# tlrl-mpla2), MF-59 (AddaVax, Cat# vac-adx-10), and AS03 (AddaS03, Cat# vac-as03-10) were purchased from InvivoGen (San Diego, CA, USA). Lipid A (Cat# 24018-s) was obtained from Peptide Institute, Inc. (Osaka, Japan), while SAS (Cat# S6322) and LPS (*E. coli* (O111:B4), Cat# L2630) were obtained from Sigma-Aldrich (St. Louis, MO, USA). Doxorubicin (Cat# 046-21523) was purchased from Sandoz K.K. (Tokyo, Japan), and TAK-242 (Resatorvid, Cat# S7455) was purchased from Selleck Chemicals (Houston, TX, USA). Spermidine (Cat# 191-13831) was purchased from FUJIFILM Wako Pure Chemical Corporation (Osaka, Japan). arvenin-1 (Cat# 65247-27) was purchased from Angene (Nanjing, China).

### Cell culture

Isolation of neonatal rat cardiomyocytes (NRCMs) was performed as described previously (30). Human iPSC-derived cardiomyocytes (hiPSC-CMs) products (iCell Cardiomyocytes 2.0) were purchased from FUJIFILM Cellular Dynamics, Inc. (Osaka, Japan) and maintained according to the manufacturer’s instructions. Samples were evaluated 20 or 24 hours after treatment with each adjuvant and compound in NRCMs and hiPSC-CMs.

### Cell viability and cytotoxicity

NRCMs (1.5 × 10^5^ cells/well) were seeded into a 96-well plate coated with Matrigel. Twenty hours after drug treatment, cytotoxicity and cell viability were evaluated using the Cytotoxicity LDH Assay Kit-WST (Dojindo Molecular Technologies, Inc., Kumamoto, Japan) and the Cell Counting Kit-8 (Dojindo Molecular Technologies, Inc., Kumamoto, Japan), respectively, in accordance with the manufacturers’ manuals. Each measured value was normalized based on the absorbance of the untreated control wells.

### Measurement of OCR

Oxygen consumption rate (OCR) was measured using an XFp Extracellular Flux Analyzer (Agilent Technologies, Santa Clara, CA, USA). NRCMs (1.5 × 10^4^ cells/well) were seeded onto Matrigel-coated XFp plates. Following the manufacturer’s protocol, 10 µM oligomycin (oligo), 2 µM FCCP, 10 µM rotenone, and 10 µM antimycin A (Rot/AA) (Mito Stress Test Kit, Cat# 103010-100, Agilent Technologies, Santa Clara, CA, USA) were administered sequentially. After OCR measurement, cells were fixed with 4% paraformaldehyde (Cat# 11850-14, Nacalai Tesque, Kyoto, Japan), washed with PBS, and stained with DAPI (Cat# 340-07971, Dojindo Molecular Technologies, Inc., Kumamoto, Japan) for 10 minutes. The number of cells in each well was measured using a BZ-X800 fluorescence microscope (Keyence, Osaka, Japan), and OCR was normalized to cell number. Human iPSC-CMs (1.0–1.5 × 104 cells/well) were seeded onto Matrigel-coated XFp plates, and OCR was measured in the same manner.

### Mitochondrial morphology

NRCMs were incubated with 200 nM MitoTracker Green FM (Cat# M46750, Invitrogen, Carlsbad, CA, USA) at 37°C for 20 minutes. After washing twice with HBSS, imaging was performed on a BZ-X800 microscope (Keyence, Osaka, Japan). Mitochondrial lengths were measured using ImageJ and classified into three categories: vesicular, intermediate, and tubular.

### Mitochondrial membrane potential and mtROS

To assess mitochondrial membrane potential, 2 µM JC-1 (Cat# MT09, Dojindo Molecular Technologies, Inc., Kumamoto, Japan) was used. After incubating at 37°C for 30 minutes, images were captured using a BZ-X800 fluorescence microscope (Keyence, Osaka, Japan). Fluorescence intensity (Red/Green ratio) was calculated with ImageJ.

For the measurement of mitochondrial ROS, 1 µM MitoSOX Red (Cat# M36008, Invitrogen, Carlsbad, CA, USA) was used. After a 10-minute incubation, the samples were washed twice with HBSS, and images were acquired using a BZ-X800 microscope (Keyence, Osaka, Japan).

### Analysis of cardiac contraction

NRCMs (1.0 × 10^6^ cells/well) were seeded into Matrigel-coated dishes. Twenty-four hours after seeding, the medium was replaced with low-glucose DMEM with 2% FBS. After washing cells with PBS, electrical stimulation was performed using platinum electrodes under the following conditions: Pulse width: 10 milliseconds; Inter-stimulus interval: 0.1 seconds; Voltage: 60 V; Frequency: 10 Hz. Starting 3 minutes after the onset of stimulation, the contraction velocity of cardiomyocytes was recorded using the SI-8000 cell motion analysis system (SONY, Tokyo, Japan) and used for analysis(31).

### Animals

All animal studies were conducted according to the guidelines concerning the care and handling of experimental animals, and approved by the Animal Care and Use Committee, Kyushu University (A23-022-2, A24-424-1, A25-164-0, and A25-309-1). Male and female 1–3 days old Sprague–Dawley rat pups for the isolation of NRCMs, and eight-week-old BALB/c mice (18–22 g, female) were purchased from Japan SLC (Shizuoka, Japan). Mice were housed in a room maintained at 21–23 °C under a 12-hour light-dark cycle, with free access to food and water. Nine-week-old mice were used for the experiments.

### Animal model

To evaluate the safety of the adjuvants, PBS, alum, AS03, MF59, SAS, and alum + lipid A were each administered intraperitoneally to BALB/c mice twice at two-week intervals. Tissue samples were collected 4 weeks after the initial administration.

To establish a heart failure model, doxorubicin (4 mg/kg) was administered intraperitoneally to BALB/c mice once a week for a total of five doses. Influenza HA (Cat# 128-213733, BIKEN, Osaka, Japan) was mixed with adjuvant (alum or alum + lipid A), and a total of 100 μL was administered subcutaneously to the mice. Each dose contained 500 ng/mouse influenza HA, 500 ng/mouse alum, and 50 ng/mouse lipid A. Vaccination was performed one week after the first doxorubicin injection, followed by a second vaccination two weeks later, for a total of two doses. To measure antibody titers, blood was collected weekly via the tail vein. Serum was recovered by centrifugation at 3,000 × g for 10 minutes. Tissue samples were collected 4 weeks after the initial vaccination. Motor function was evaluated using treadmill (Cat# TMS-M2, MELQUEST, Toyama, Japan).

### Analysis of cardiac function

Cardiac function in mice was measured weekly under anesthesia (1–2% isoflurane) using echocardiography: Prospect T1 (S-Sharp Corporation, New Taipei, Taiwan). Left ventricular ejection fraction (LVEF) and left ventricular fraction of shortening (LVFS) were calculated using two-dimensional M-mode imaging of the left ventricular long-axis view. We calculated the average value of three consecutive heart cycles.

### ELISA

HA protein antigen (100 ng/well) diluted in Coating Buffer (27.6 mM Na2CO3, and 3.08 mM NaN3) was dispensed into a 96-well plate and incubated overnight at 4°C. After washing with PBST (PBS containing 0.05% Tween-20), PBST containing 10% FBS was added as a blocking solution and incubated at room temperature for 1 hour. Collected mouse serum (100 μL/well) diluted 50,000-fold in the blocking solution was added, and the mixture was incubated at room temperature for 2 hours. After washing, HRP-labeled secondary antibody (Goat F(ab) Anti-Mouse IgG (HRP), Cat# ab6833, Abcam, Tokyo, Japan) was diluted 25,000-fold in PBST and incubated at room temperature for 1 hour. After washing, TMB substrate solution (Cat# 00-4201-56, Invitrogen, Vienna, Austria) was added and incubated at room temperature for 20 minutes, then the reaction was stopped with H2SO4 solution. The absorbance of each well was measured using Nivo (PerkinElmer, Waltham, MA, USA), and the data were used for analysis.

### Immunohistochemistry

Spleen tissue was embedded in O.C.T. compound (Cat# 90501, Sakura Finetek, Torrance, CA, USA) and rapidly frozen. Frozen sections 12 μm thick were prepared using a Cryostat HM525 NX (Thermo Fisher Scientific, Waltham, MA, USA). Tissue slices were washed with PBS, PBST containing 10% FBS was added as a blocking solution, and incubated at room temperature for 1 hour. PE-labeled anti-mouse CD4 antibody (Cat# 130310, BioLegend, San Diego, CA, USA) and Alexa Fluor 700-labeled anti-mouse/human CD45R/B220 antibody (Cat# 103231, BioLegend, San Diego, CA, USA) were added and incubated overnight at 4°C. After washing with PBST, Alexa Fluor 568-labeled goat anti-rat IgG secondary antibody (Cat# 2379471, Invitrogen, Carlsbad, CA, USA) and DAPI (Cat# 340-07971, Dojindo Molecular Technologies, Inc., Kumamoto, Japan) were added, and the samples were incubated at room temperature for 1 hour. After washing, the samples were sealed using FluoroKeeper (Cat# 12593-64, Nacalai Tesque, Kyoto, Japan). Images were obtained using a BZ-X800 microscope (Keyence, Osaka, Japan), and the areas of CD4- and CD45R/B220-positive regions in each spleen region were calculated using ImageJ software (V1.54g, National Institutes of Health, Bethesda, Maryland, USA).

### Statistical analysis

All results are presented as the mean ± S.E.M. Statistical significance was assessed using a one-way analysis of variance (ANOVA) followed by Tukey’s multiple comparison test (when there were three or more groups), and a two-way analysis of variance (ANOVA) followed by Bonferroni’s multiple comparison test (when there were three or more groups with variables that changed over time). P-value of less than 0.05 was considered statistically significant.

## Results

### Adjuvants do not affect cardiac function and cell viability

To confirm the toxicity of the adjuvants used in this study, we evaluated them using mice and NRCMs. There were no significant differences in LVEF and LVFS between mice administered the five types of adjuvants and control mice after 4 weeks (Fig. 1A, B). Regardless of whether the adjuvant was administered, there were no changes in body weight or heart weight (Fig. 1C, D). Adjuvants did not affect either the cytotoxicity or cell viability of NRCMs (Fig. 1E, F). These results reveal that the adjuvants do not affect the heart.

**Figure 1:**
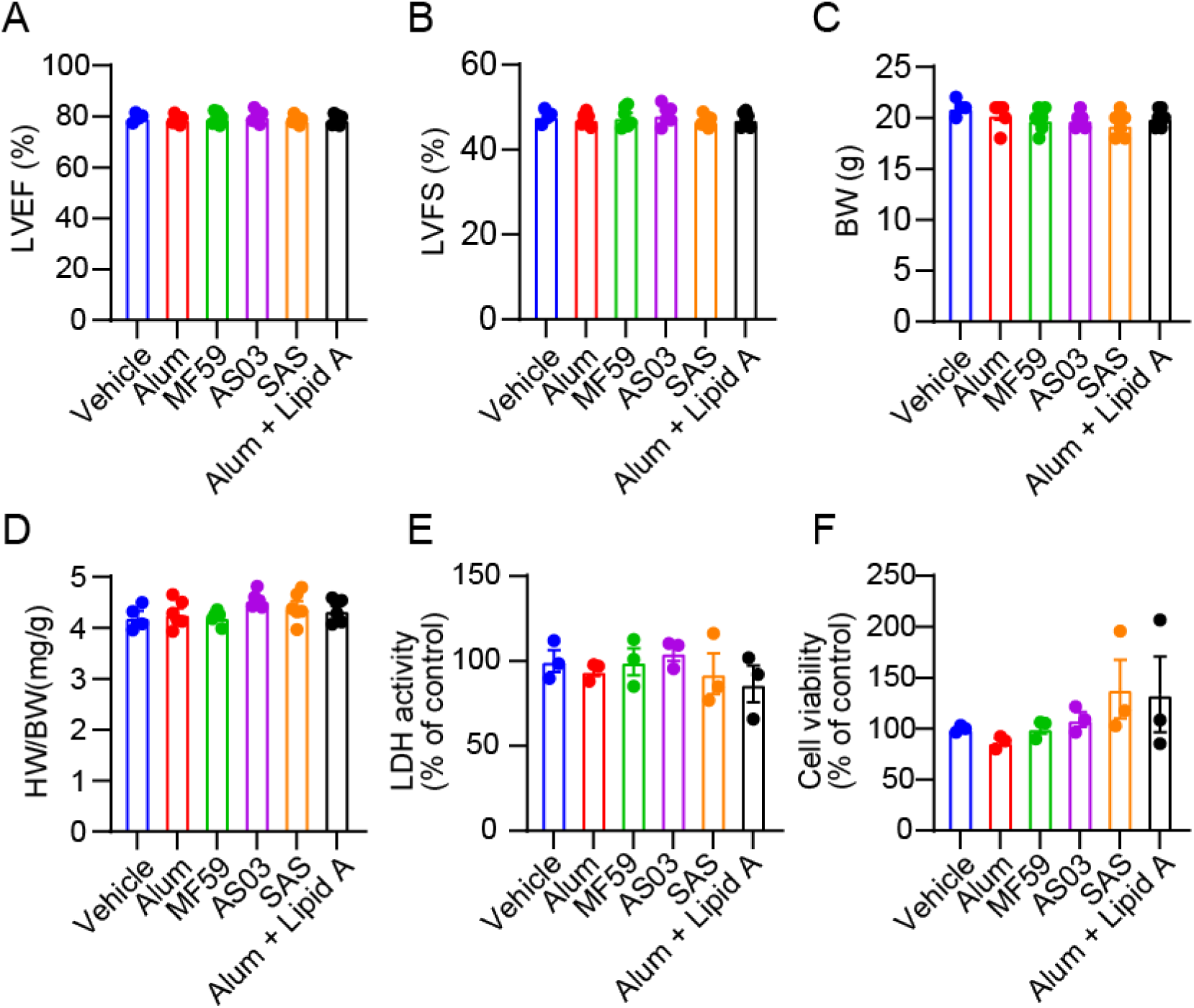
These adjuvants do not affect cardiac function. (A, B) LVEF (A) and LVFS (B) 4 weeks after adjuvant administration. (C, D) Body weight (C) and heart weight (D) 4 weeks after adjuvant administration (n = 4-6 mice per group). (E, F) Cytotoxicity in NRCMs 20 hours after adjuvant treatment. Cell viability and toxicity were measured by LDH release and MTT assay (n = 3). All data are shown as mean ± SEM; Data were analyzed using one-way ANOVA followed by Tukey’s comparison test.

### Lipid A enhances myocardial mitochondrial function

Next, we evaluated the effects of the adjuvant on myocardial mitochondria. Most of the adjuvants did not affect respiratory capacity; however, maximal respiration was significantly increased only in the alum + lipid A group (Fig. 2A). Following treatment of NRCMs with lipid A, maximal respiratory capacity increased in the group treated with lipid A (Fig. 2B, C). Treatment with lipid A did not alter the morphology of cardiac mitochondria (Fig. 2D, E). Mitochondrial membrane potential (Δψm) was evaluated using JC-1 staining. Lipid A induced the hyperpolarization of the mitochondrial membrane potential (Fig. 2F, G). On the other hand, alum did not affect mitochondrial OCR or morphology (Fig. 2H-K).

**Figure 2:**
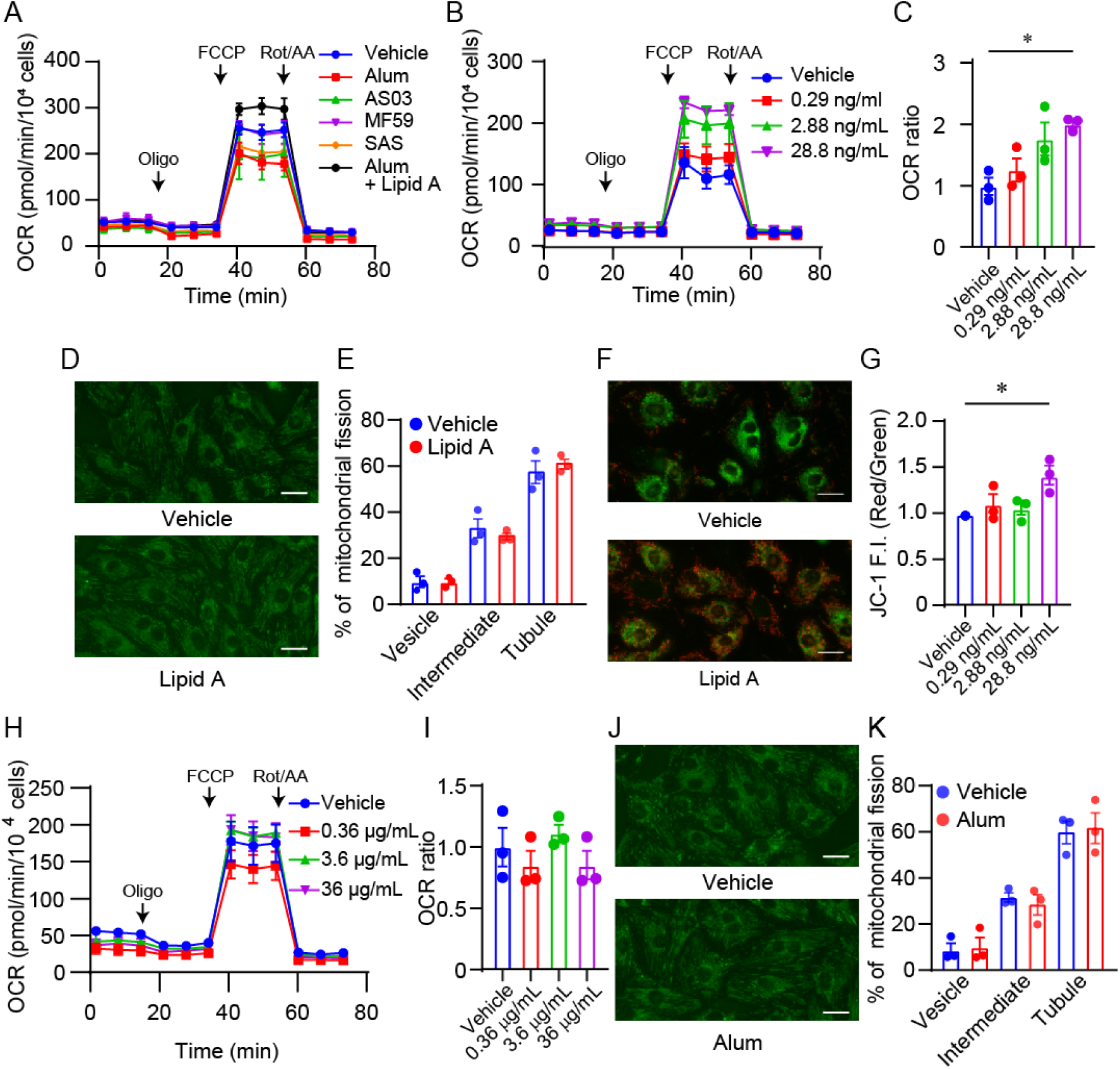
Lipid A enhances myocardial mitochondrial function. (A, B, H) OCR in NRCMs 20 hours after treatment with various adjuvants (A), lipid A (B), and alum (H). (C, I) Maximal respiratory rate in NRCMs 20 hours after treatment with lipid A (C), and alum (I). (D, J) Representative images of mitochondria in cardiomyocytes treated with lipid A (D) and alum (J) were stained by MitoTracker Green FM. (E, K) Comparison of mitochondrial morphology in cardiomyocytes in lipid A (E) and alum (K)-treated group. The cells were divided into three groups (Vesicle, Intermediate, Tubule) based on the length of mitochondria per cell in alum or lipid A treatment groups. (F) Representative images of mitochondrial membrane potential after treatment with lipid A (G) The ratio of red fluorescence intensity to green fluorescence intensity was quantified and normalized to the vehicle group. Scale bar: 20 μm. All data are shown as mean ± SEM; n = 3. *P < 0.05. Data were analyzed using one-way ANOVA followed by Tukey’s comparison test.

### Lipid A regulates myocardial mitochondrial respiration through TLR4

Lipid A is recognized as the center of the immune response in lipopolysaccharides (LPS) derived from Gram-negative bacteria. It is widely known to elicit an immune response via Toll-like receptor 4 (TLR4) on the cell surface. TAK-242, a TLR4-specific inhibitor, suppressed the enhancement of myocardial mitochondrial respiratory capacity induced by lipid A (Fig. 3A, B). Treatment with the TLR4 agonists MPL or LPS did not result in any changes in mitochondrial respiratory capacity or mitochondrial membrane potential (Fig. 3C-F, SFig. 1A-D). These results suggest that while lipid A targets TLR4, it induces activation mechanisms distinct from those of MPL and LPS.

**Figure 3:**
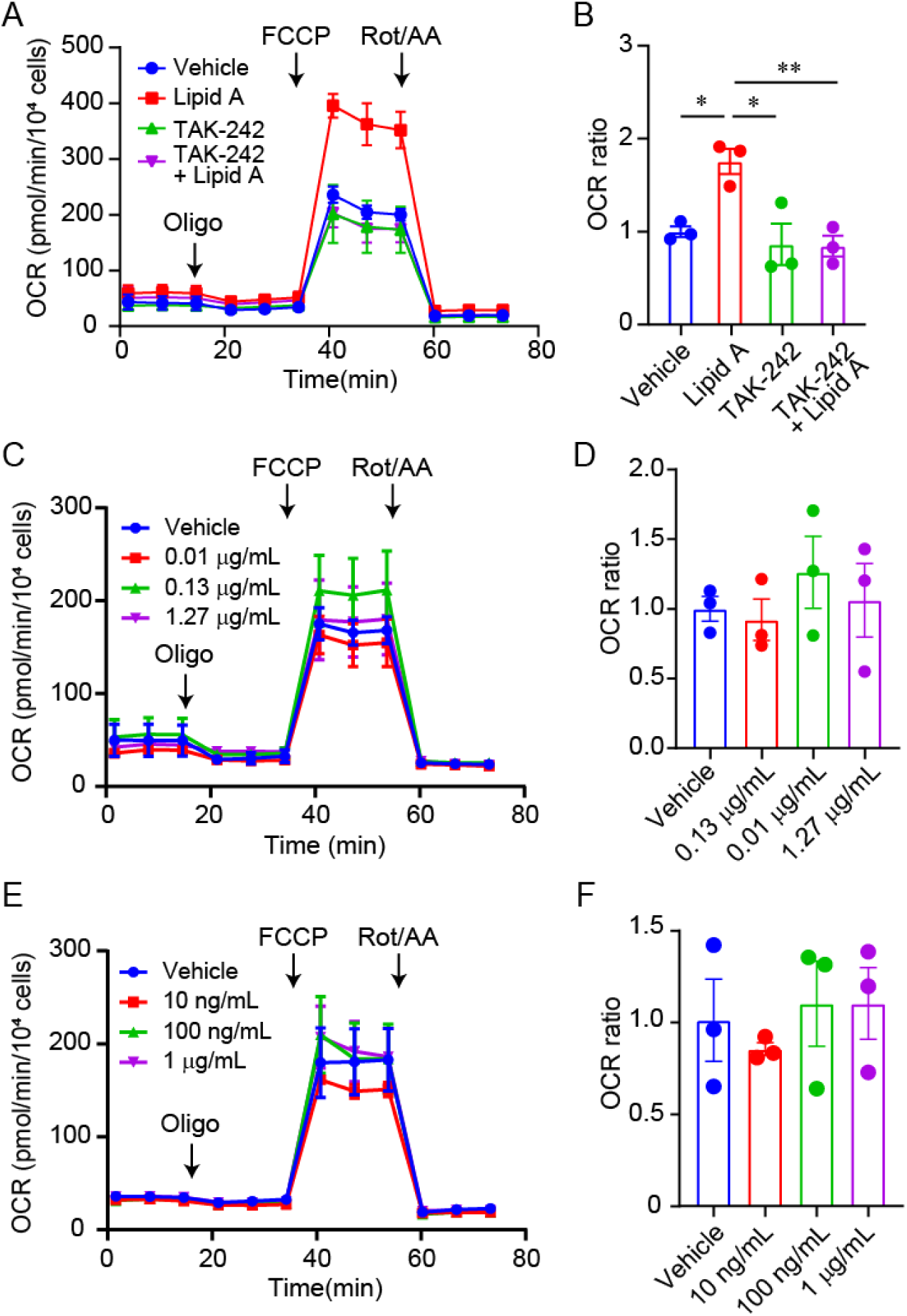
Lipid A regulates myocardial mitochondrial respiration via TLR4. (A) OCR in NRCMs treated with lipid A for 24 hours following 1-hour pretreatment with TAK-242. (C, E) OCR in NRCMs 24 hours after treatment with MPL (C), and LPS (E). (B, D, F) Maximal respiratory rate in each condition (A, C, E). All data are shown as mean ± SEM; n = 3. *P < 0.05, **P < 0.01. Data were analyzed using one-way ANOVA followed by Tukey’s comparison test.

### Lipid A suppresses doxorubicin-induced mitochondrial dysfunction in cardiomyocytes

Doxorubicin (Dox), a potent anticancer drug, exhibits excellent antitumor effects. However, as a side effect, it induces concentration-dependent cardiotoxicity. It has been reported that Dox significantly impairs myocardial mitochondrial respiratory capacity (16). Lipid A suppressed Dox-induced mitochondrial dysfunction (Fig. 4A, B). We treated cardiomyocytes with spermidine and arvenine-1, which have been reported to restore antitumor immunity by activating mitochondria in immune cells. Although spermidine and arvenine-1failed to enhance mitochondrial function, arvenine-1 showed a trend toward increased maximal respiratory capacity (SFig. 2A-D). Dox treatment significantly reduced the contractility of NRCMs, but this contractile impairment was improved by lipid A treatment (Fig. 4C, D). Furthermore, we evaluated the production of ROS induced by Dox (Fig. 4G, H). As with NRCMs, Dox treatment reduced OCR in hiPSC-CMs; however, this reduction was significantly ameliorated by the administration of lipid A (Fig. 4E, F). While Dox treatment significantly increased mitochondrial ROS production, lipid A treatment significantly suppressed this excessive ROS production (Fig. 4G, H). These results demonstrated that lipid A suppresses Dox-induced mitochondrial dysfunction and myocardial contractile impairment.

**Figure 4:**
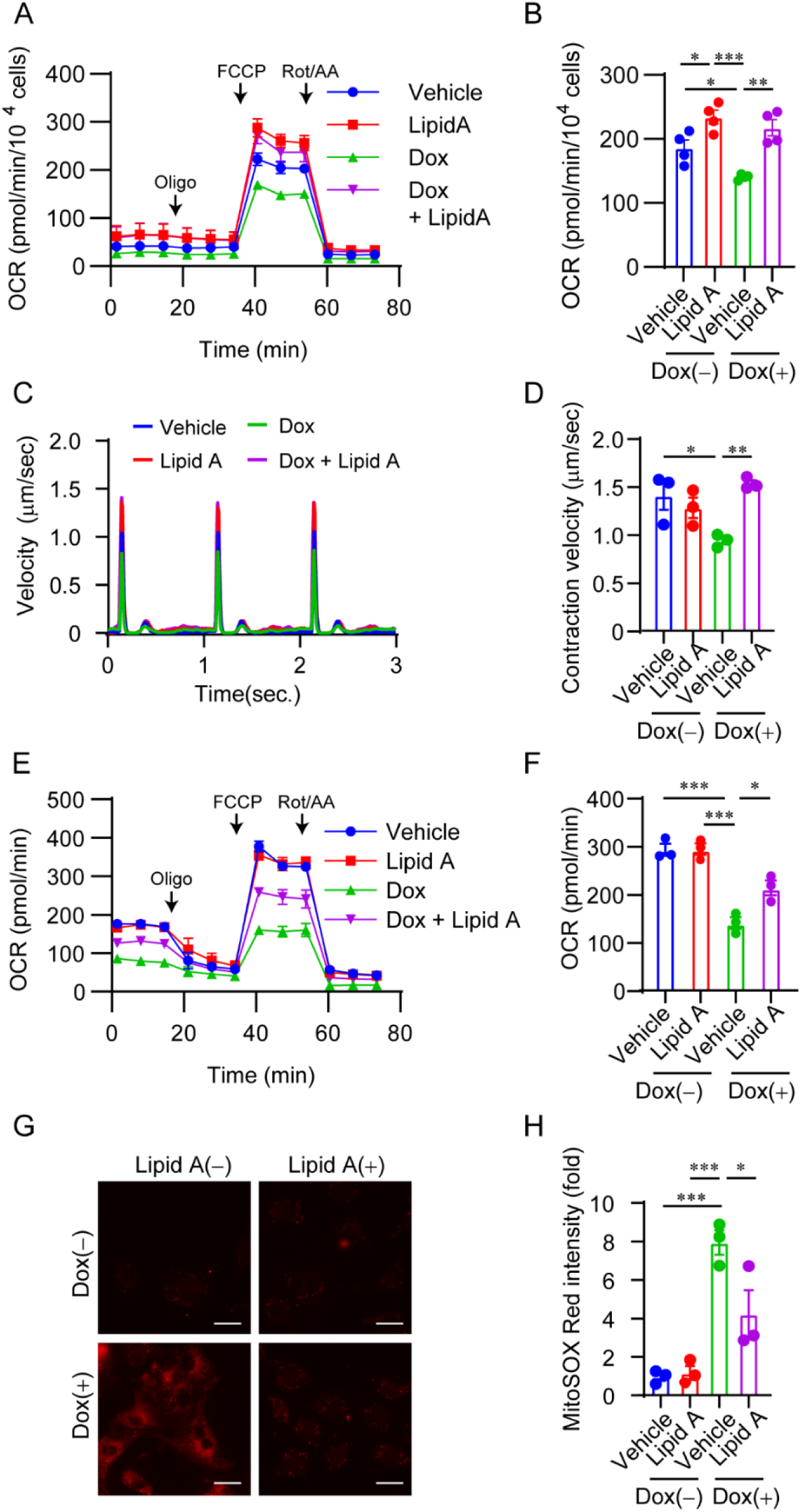
Lipid A suppresses doxorubicin-induced mitochondrial dysfunction in rat and human cardiomyocytes. (A) OCR in NRCMs 24 hours after treatment with 50 nM Doxorubicin (Dox), and lipid A. (B) Maximal respiratory rate in (A). (C, D) Representative traces (C) and quantification of contractile velocity (D) in NRCMs 24 hours after treatment with 50 nM Dox and lipid A. (E) OCR in hiPSC-CMs 24 hours after treatment with 1 μM Dox, and lipid A. (F) Maximal respiratory rate in (E). (G) Representative images (G) of mitochondrial ROS 24 hours after treatment with 100 nM Dox and lipid A. (H) MitoSOX Red fluorescence intensity was measured and normalized to the vehicle group without Dox as a control. Scale bar: 20 μm. All data are shown as mean ± SEM; (A, B) n = 4, (C-H) n = 3.*P < 0.05, **P < 0.01, ***P < 0.001. Data were analyzed using one-way ANOVA followed by Tukey’s comparison test.

### Lipid A improves doxorubicin-induced cardiotoxicity and immunosuppression

Dox is known to cause severe cardiotoxicity and immunosuppression in a dose-dependent manner. Therefore, vaccination is considered important in practice to reduce the risk of infections in cancer patients. Next, we administered a vaccine with lipid A to Dox-induced heart failure model mice and evaluated its effects. Using alum, which is commonly used in clinical practice, as a control, a mixture of alum and lipid A was used as an adjuvant, and influenza HA was used as the antigen (Fig. 5A). Cardiac function began to decline gradually one week after Dox administration and had significantly deteriorated by two weeks (Fig. 5B, C). The significant decreases in LVEF and LVFS observed in the Dox-treated group were significantly attenuated by the administration of alum and lipid A (Fig. 5B, C). The decrease in heart weight and running distance caused by Dox was improved by lipid A treatment (Fig. 5D, E). In Dox-treated mice administered alum, serum antibody titers decreased significantly; however, this decrease was suppressed by the concurrent administration of lipid A (Fig. 5F). The reduction in spleen weight caused by Dox was also alleviated by lipid A treatment (Fig. 5G). To elucidate the mechanisms underlying immune function maintenance, we stained these spleens using CD4 (a T cell marker) and CD45R/B220 (a B cell marker). No significant changes in the area of CD4-positive regions were observed between groups (Fig. 5H, I). In contrast, the area of CD45R/B220-positive regions decreased with Dox, but lipid A suppressed this decrease in area (Fig. 5J, K). These results indicated that lipid A attenuates cardiotoxicity by maintaining myocardial mitochondrial function and protecting the B cell region in the spleen, thereby sustaining the immune response to the vaccine even under doxorubicin administration.

**Figure 5:**
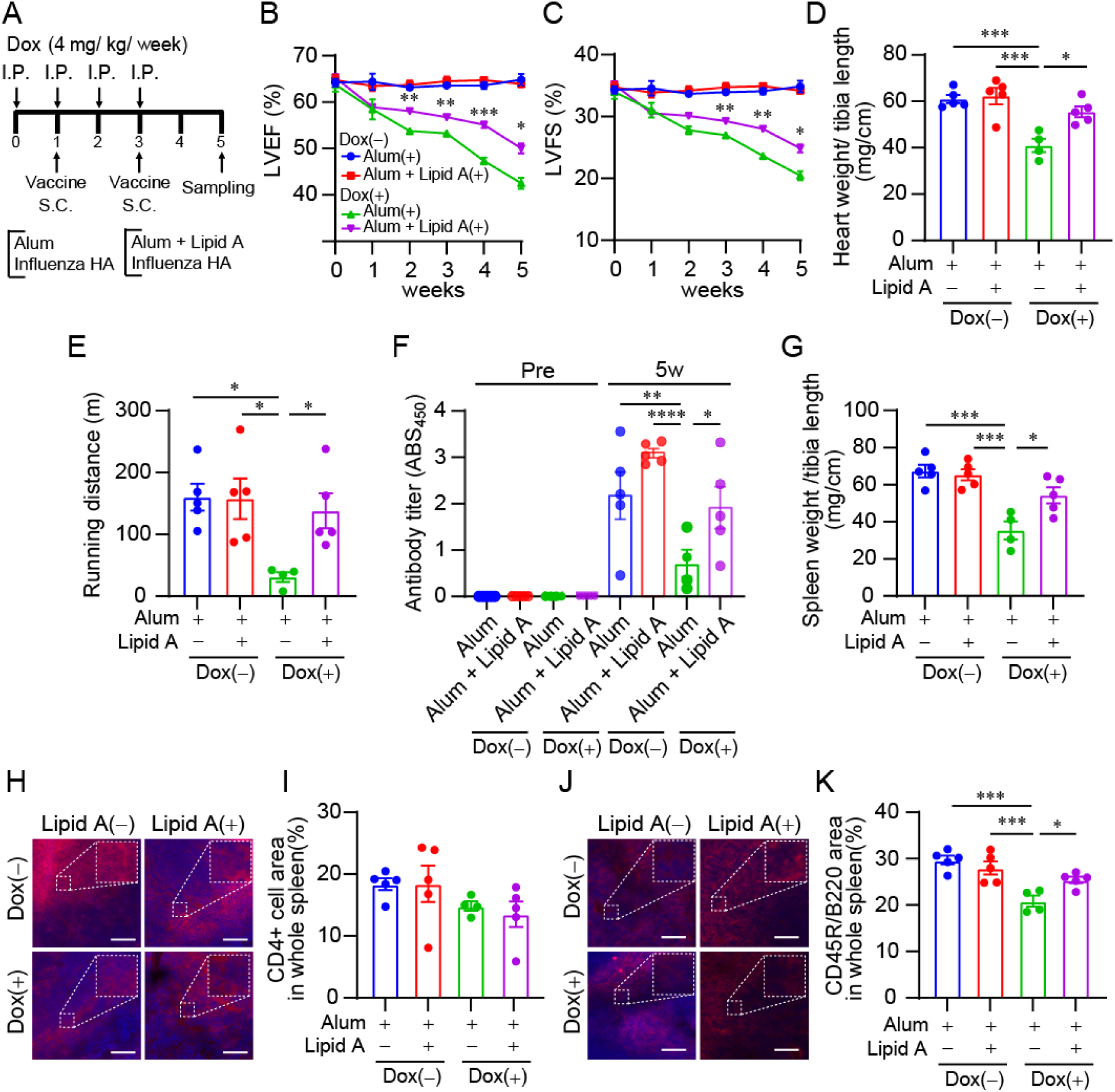
Lipid A improves doxorubicin-induced cardiotoxicity and immunosuppression. (A) Experimental protocol. The first vaccine dose was administered one week after doxorubicin administration, followed by second dose two weeks later. Influenza HA was used as the antigen, and two adjuvant groups were established: a control group receiving only alum and a group receiving alum and lipid A. Cardiac function was measured weekly. (B, C) LVEF (B) and LVFS (C) after Dox treatment in time-dependent manner. (D, E) Heart weight/tibia length ratio (D) and running distance (E) in mice 5 weeks after Dox treatment. (F) Serum antibody titer against influenza HA at the start of the experiments and at week 5. (G) Spleen weight/tibia length ratio in mice 5 weeks after Dox treatment. (H, J) Representative images of spleen by IHC staining. CD4-positive cells (H) and CD45R/B220-positive cells (J) are shown, with nuclei stained with DAPI (blue). (I, K) Quantification of the area of CD4-positive cells (I) and CD45R/B220-positive cells (K) in the spleen. Scale bar: 200 μm. All data are shown as mean ± SEM; n = 4-5. *P < 0.05, **P < 0.01, ***P < 0.001, ****P < 0.0001. Data were analyzed using one-way ANOVA followed by Tukey’s comparison test, and a two-way ANOVA followed by Bonferroni’s multiple comparison test.

## Discussion

This study evaluated the effects of various adjuvants on the heart, focusing on myocardial mitochondria. Among the five adjuvants tested, lipid A uniquely enhanced the maximal respiratory capacity of myocardial mitochondria. Furthermore, we demonstrated that lipid A improves mitochondrial dysfunction of cardiomyocytes and ameliorates impaired cardiac function induced by the anticancer drug Dox.

Administration of alum, MF59, AS03, SAS, and lipid A to wild-type mice did not change cardiac function or body and heart weight (Fig.1 A-D). When these adjuvants were administered to NRCMs, no cytotoxicity was observed compared to the vehicle group, indicating that these adjuvants do not cause cardiac dysfunction under physiological conditions (Fig.1 E, F).

Of the five adjuvants, only lipid A enhanced maximal mitochondrial respiratory capacity (Fig. 2A). Although lipid A did not affect mitochondrial morphology, it also induced an increased mitochondrial membrane potential (Fig. 2B, C). Since proton transport from the inner mitochondrial membrane to the outer membrane by Complexes I, III, and IV is critical for mitochondrial membrane potential generation, these results suggest that lipid A may enhance the activity of electron transport chain.

Previous reports have shown that lipid A activates TLR4 (32). Treatment with TAK-242, an inhibitor of TLR4, suppressed the increase in maximal respiration induced by lipid A, indicating that TLR4 is involved in this mechanism of action (Fig. 3A). Next, treatment of NRCMs with the TLR4 agonists MPL and LPS did not increase maximal respiration (Fig. 3B, C). These findings indicate that there are differences in downstream signaling pathways between lipid A and MPL/LPS-mediated TLR4 activation. LPS binds to MD-2 hydrophobic pocket and activates intracellular signaling by mediating the dimerization of TLR4-MD-2 complexes (33). Subsequently, it activates NF-κB, induces increased release of inflammatory cytokines and mitochondrial oxidative stress, and causes mitochondrial dysfunction (34, 35). Recently, it has been reported that LPS-mediated TLR4 activation in skeletal muscle promotes excessive mitochondrial fission (36). In BMDCs, it has been reported that while both MPL and lipid A target TLR4, their downstream signaling pathways differ (32). Unlike MPL, lipid A is dependent on myeloid differentiation factor 88 (MyD88) signaling (37, 38). Furthermore, it has been reported that MF59 functions via the MyD88 activation pathway in a TLR-independent manner (39). However, since MF59 does not affect mitochondrial function (Fig. 2A), this suggests that the TLR4-MyD88 pathway contributes to the enhancement of mitochondrial respiratory capacity. Although both LPS and lipid A are known to activate the TLR4–MyD88 pathway, the reason for their distinct effects on mitochondrial function remains unclear.

Since lipid A enhanced mitochondrial function, we used a Dox model that impairs mitochondrial function to verify its therapeutic effects. The maximal respiratory capacity, which was suppressed by Dox, was restored by lipid A not only in NRCMs but also in hiPSC-CMs (Fig. 4A, E). Furthermore, myocardial contractility was also improved in NRCMs (Fig. 4C). These effects are thought to result from lipid A protecting mitochondria from the excessive oxidative stress induced by Dox. Next, we examined the effects of lipid A on not only cardiac function but also immune activation in a Dox-induced heart failure mouse model. Various mechanisms have been reported to contribute to Dox-induced cardiotoxicity, including oxidative stress, mitochondrial dysfunction, activation of apoptotic pathways, and endoplasmic reticulum (ER) stress (21–23). Lipid A significantly suppressed Dox-induced cardiac dysfunction and improved the decline in exercise capacity associated with this dysfunction (Fig. 5B–E). These results suggest that lipid A acts on mitochondria to suppress Dox-induced cardiotoxicity. Furthermore, while anti-influenza HA antibody titer was reduced in Dox-treated mice, high antibody titers were maintained in the group treated with lipid A (Fig. 5F). Analysis of the spleen phenotype related to antibody production revealed that lipid A suppressed the reduction in spleen weight (Fig. 5G). In the spleen, there was no difference in the proportion of T cells (CD4^+^), among the groups; however, the proportion of B cells (CD45R/B220^+^) decreased following Dox treatment but was restored by lipid A administration. It is speculated that Dox induced apoptosis in B cells, and the suppression of the decrease in B cells by lipid A led to the persistence of antibody titers.

Currently, vaccination during chemotherapy is recommended in clinical guidelines; however, the depletion of blood cells caused by bone marrow suppression resulting from anticancer drug therapy, as well as the reduced efficacy of vaccines, remains a significant challenge (26, 40–42). Furthermore, mitochondrial metabolism is crucial for the proliferation and differentiation of immune cells, and mitochondrial dysfunction caused by the tumor microenvironment or metabolic stress leads to a reduced response in APCs and lymphocytes (43–46). Lipid A has the potential to address these challenges. This study demonstrated that lipid A does not affect cardiac function and, unlike conventional adjuvants, not only activates the immune system but also enhances mitochondrial function. In addition, it suppressed Dox-induced cardiotoxicity. The fact that an adjuvant can produce multiple pharmacological effects may provide innovative insights for future adjuvant development.

## Acknowledgment

We sincerely thank Ms. Yukina Ishii for preliminary experiments. This work was supported by JSPS KAKENHI under Grant Numbers 23K06164 (to Yu.K.), 26H02407 (to M.N.) from the Ministry of Education, Culture, Sports, Science and Technology of Japan, AMED SCARDA under Grant Numbers JP223fa727001 (to M.U., Ya.K., J.K., and M.N.), and JP223fa827001 (to M.N.), AMED under Grant Numbers JP24mk0121280 (to Ya.K. and Yu.K.), JP25ama121031 (to M.N.), JP26gm1910010 (to M.N.), the Smoking Research Foundation (to M.N.), and the Fukuda Foundation for Medical Technology (to Yu.K.).

