## Supplemental Figure for "Lipid A counteracts doxorubicin-induced systemic dysfunction by boosting mitochondrial activity": BioRxiv Supplementary Figure (Nakaguma et al) 260417★.pdf

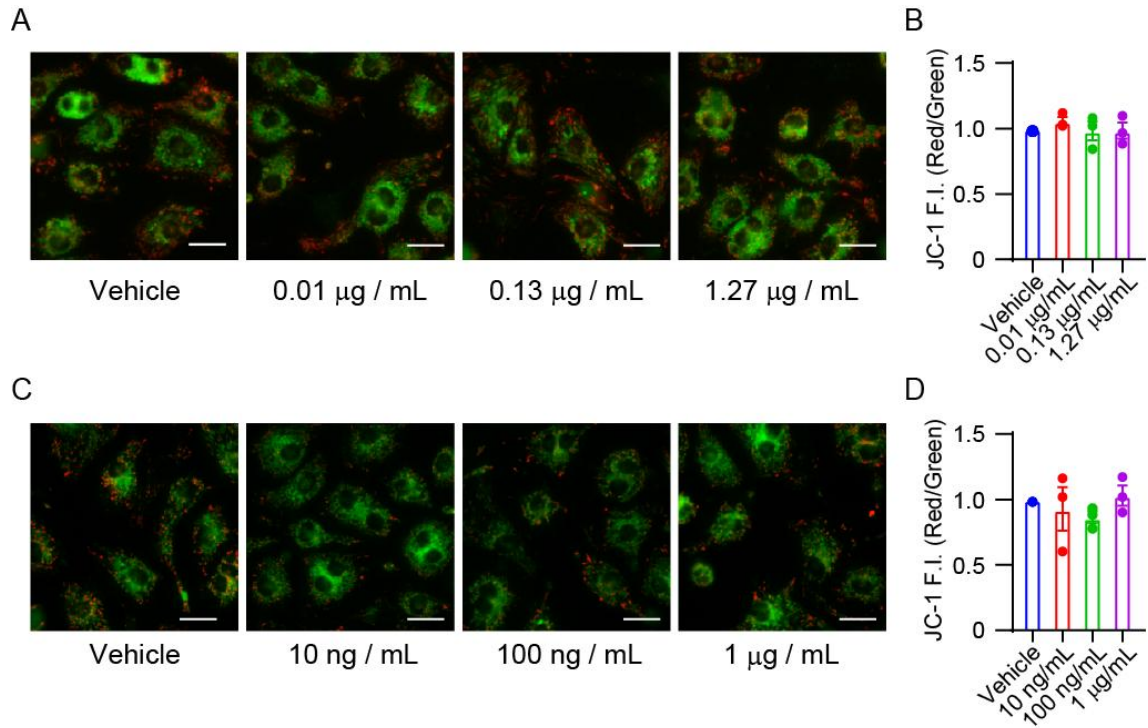

**Supplementary Figure 1: MPL and LPS fail to hyperpolarize mitochondrial membrane potential.**

(A-D) Representative images of mitochondrial membrane potential 20 hours after treatment with MPL (A, B) and LPS (C, D). The ratio of red fluorescence intensity to green fluorescence intensity was quantified and normalized to the vehicle group. All data are shown as mean ± SEM; $n = 3$. Data were analyzed using one-way ANOVA followed by Tukey's comparison test.

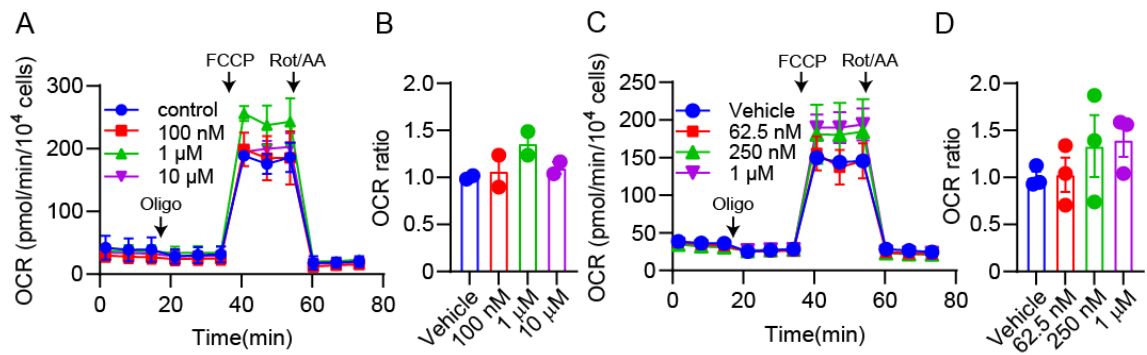

**Supplementary Figure 2: Spermidine and arvenin-1 fail to enhance myocardial mitochondrial function.**

(A-D) OCR in NRCMs 20 hours after treatment with spermidine (A), arvenin-1 (C). (B, D) Maximal respiratory rate after treatment with spermidine (B), and arvenin-1 (D). All data are shown as mean ± SEM; $n = 3$. Data were analyzed using one-way ANOVA followed by Tukey's comparison test.

data are shown as mean ± SEM; n = 3. Data were analyzed using one-way ANOVA followed by Tukey's comparison test.
